## Supplementary Material for "Optimized Vivid-derived Magnets photodimerizers for subcellular optogenetics"

#### SUPPLEMENTAL INFORMATION

#### SUPPLEMENTARY FIGURE LEGENDS

**Supp. Figure 1: Domain organization diagrams of the constructs used in this study.**

A. Constructs used to induce wild-type and mutant Magnets heterodimerization at the outer mitochondrial membrane.

B. Bait proteins used for the light-dependent recruitment of soluble prey proteins to the ER and lysosomes.

C. Vectors used for the light-dependent modulation of $PI(4,5)P_2$ at the plasma membrane.

D. Constructs used for the optogenetic induction of organelle contacts.

E. Constructs used to induce VAP reconstitution at the surface at the ER membrane (Opto-VAP). MSP, major sperm protein homology domain; CC, coiled-coil domain; TM, transmembrane domain; PH, Pleckstrin homology domain; FFAT, two phenylalanines (FF) in an Acidic Tract motif; ORD, OSBP-related protein lipid-binding domain.

**Supp. Figure 2: Recruitment of the cytosolic prey to the membrane-associated bait upon light stimulation**

Accumulation of a soluble prey (eMagB$_F$-TagRFP-T or eMagB-TagRFP-T) to a mitochondria-anchored bait (eMagA$_F$-EGFP-Mito) (A) or to an ER-associated bait (ER-EGFP-eMagA) (B) upon whole-cell illumination in a HeLa cell (A) or COS-7 cell (B). Scale bar: 2 μm.

**Supp. Figure 3: Magnets mutations tested to improve heterodimerization efficiency and thermodynamic stability.**

A. Primary sequence of the *Neurospora crassa* (strain ATCC 24698) photoreceptor Vivid (UniProtKB Q1K5Y8_NEUCR). The N-cap dimerization domain, the Per-ARNT-Sim (PAS) core domain (the photosensitive portion of the protein), and the amino acids involved in binding the flavin adenine dinucleotide (FAD) cofactor (Heintzen et al., 2001; Zoltowski et al., 2007) are indicated. The construct used for the crystal structure of the homodimer (see below) lacks the N-terminal 36 a.a.; Magnets also lack these 36 a.a. Arrows point to a.a. mutated in an attempt to generate Magnets with improved charge complementarity (see Table in panel D). Amino acid substitutions introduced during the screen are shown below the WT sequence. Magenta:

mutations that abolish potential ubiquitination sites; blue, mutations predicted to improve packing or secondary-structure preference; orange, mutations that mimic corresponding residues in thermophilic ascomycetes. The effects on heterodimerization of each substitution and combinations of substitutions is summarized in Supplementary Table 3.
B. Crystal structure of the Vivid homodimer and domain cartoon of the monomer. The two monomers are shown in orange and pink. Residues 52Ile and 55Met are shown as spheres.
C. Magnets were generated by introducing mutations to make the interface of one Vivid negatively charged (Ile52Asp/Met55Gly, "negative Magnet"), and the interface of the other Vivid positively charged (Ile52Arg/Met55Arg, "positive Magnets"), so that blue light radiation leads to hetero- rather than homo-dimerization. Residues D52/G55 in the "negative Magnet" and R52/R55 of the "positive Magnet" are shown as spheres.
D. Amino acid mutations tested for optimizing charge complementarity. (Scores: (-) less efficient than the original photoreceptors, (+) more efficient, (++) much more efficient).

**Supp. Figure 4: Alignment of Vivid domain sequences from thermophilic ascomycetes.** Sequences were retrieved from NCBI. The two *Rhizomucor* sequences come from incomplete whole-genome sequencing projects. The nine mutations in eMags are shown in bold; all but one mutation was to an amino acid occurring in one of the thermophilic homologues. Surprisingly, the 55Ala mutation introduced by rational design was subsequently found to occur at that position in many thermophilic homologues.

**Supp. Figure 5: Molecular modeling of effects of specific eMags mutations.**
A. Wild-type Thr69 leaves an unsatisfied hydrogen bond donor and acceptor at the dimer interface and packs poorly. Thr also prefers strand over helix (this position is in a helical turn).
B. Thr69Leu shows much greater hydrophobic packing at the dimer interface, including with Leu69 on the other dimer half. Leu also prefers helix over strand.
C. Wild-type Met179 is a sub-standard hydrophobic packer and has high residue entropy. Met also has only modest preference for strand over helix.
D. Met179Ile makes more hydrophobic contacts and has low residue entropy. Ile also has excellent preference for strand over helix.
E. Wild-type Ser99 makes a weak "helix N-cap" hydrogen bond to the backbone amide H of

residue 102. Ser is also only weakly preferred as a residue in a helix N-cap turn.
F. Ser99Asn makes a strong "helix N-cap" hydrogen bond to the backbone amide H of residue 102. Asn is also by far the most strongly preferred residue in a helix N-cap turn.
G. Wild-type Arg136 makes an ion pair with a phosphate of FAD, but quite closely approaches the adenine ring, with which it has suboptimal packing and electrostatics.
H. Arg136Lys still makes the ion pair with a phosphate of FAD, and less closely approaches the adenine ring, decreasing unfavorable contacts.

**Supp. Figure 6: Light-dependent heterodimerization of the original Magnets at the mitochondrial surface with or without preincubation of cells at 28°C.**
A. Representative example of the light-dependent recruitment of the original Magnets prey (pMagFast2-TagRFP-T) to mitochondria in HeLa cells expressing the mitochondrial bait nMagHigh1-EGFP-Mito without (top) or with (bottom) a preincubation at 28°C. Scale bar: 5 μm.
B. Plot showing the accumulation of soluble prey from the cytosol to mitochondria in cells expressing the original Magnets either without or with a preincubation at 28°C for 12-24 hours prior to imaging and irradiation, as shown in the schematic at left (N = original Magnets: 12 cells, original Magnets (28°C): 17 cells; from 3 independent experiments).

**Supp. Figure 7: Opto-VAP reconstitution induces PI4P loss from the Golgi complex and this effect is blocked by ITZ treatment.**
The same wild-type HeLa cell expressing TagRFP-T-MSP(VAPB$_{(1\text{-}218)}$)-eMagB, ER-EGFP-eMagA and the PI4P reporter iRFP-P4C was imaged before (A) and 45 min after ITZ treatment (B). Before ITZ treatment, we could detect a reduction in iRFP-P4C (PI4P) from the Golgi upon illumination (panel A, lower row). However, after ITZ incubation, we could not visualize iRFP-P4C loss upon illumination despite efficient activation of Opto-VAP (panel B, lower row). Scale bar: 5 μm.

**Supp. Figure 8: ITZ treatment blocks PI4P loss from the Golgi complex but does not affect Opto-VAP reconstitution in WT and VAP DKO HeLa cells.**

Wild-type (A) and VAP-DKO (B) HeLa cells expressing TagRFP-T-MSP($VAPB_{(1-218)}$)-eMagB, ER-EGFP-eMagA and the PI4P reporter iRFP-P4C imaged 30 min after ITZ treatment. Despite the rapid and efficient association of TagRFP-T-MSP($VAPB_{(1-218)}$)-eMagB to ER membranes upon light stimulation (graphs at right), ITZ prevented iRFP-P4C (PI4P) loss from the Golgi in wild-type HeLa cells and endosome/Golgi hybrid structures in HeLa VAP-DKO cells. Scale bar: 5 µm. (HeLa WT: $\tau_{ON}$ = 50.7 ± 3.2 s, N=24; HeLa WT + ITZ: $\tau_{ON}$ = 46.5 ± 2.7 s, N=16; HeLa VAP-DKO: $\tau_{ON}$ = 81.1 ± 9.6 s, N=20; HeLa VAP-DKO + ITZ: $\tau_{ON}$ = 68.1 ± 8.2 s, N=16; from 3 independent experiments).

**Supp. Figure 9: $PH_{OSBP}$ mediated tethering between the ER and PI4P-rich subcellular membranes is not associated with PI4P loss from these membranes**

A. Graphical representation of the assay used to mediate VAP-independent membrane tethering of PI4P-enriched Golgi and endosomal membranes to the ER. Cells were transfected with 1) the PH domain of OSBP fused to TagRFP-T and eMagB (TagRFP-T-eMagB-$PH_{OSBP}$), 2) an ER bait (ER-EGFP-eMagA), and 3) a PI4P reporter (iRFP-P4C). Before light activation (Dark), TagRFP-T-eMagB-$PH_{OSBP}$ is partially cytosolic and already partially bound to PI4P-enriched Golgi membranes (in WT cells) and Golgi/endosome hybrid membranes (in VAP-DKO cells). Upon blue-light illumination, eMags associated with the ER, including ER in proximity of the Golgi and endosomes, brings PI4P-enriched Golgi, or hybrid endosome-Golgi organelles, in close apposition to the ER, but no PI4P loss occurs given the absence of the ORD.

B. High-magnification view of the Golgi complex of a WT cell and of the hybrid Golgi/endosome organelles of a VAP-DKO HeLa cell expressing the constructs indicated in (A). The iRFP-P4C (PI4P) signal is shown. Blue light-dependent formation of the tether does not reduce PI4P levels on these organelles. Scale bar: 5 µm (WT) and 2 µm (DKO).

C. Quantification of the results shown in (B). WT HeLa (N= 16); VAP-DKO HeLa (N= 17).

### Supplementary Figure 1

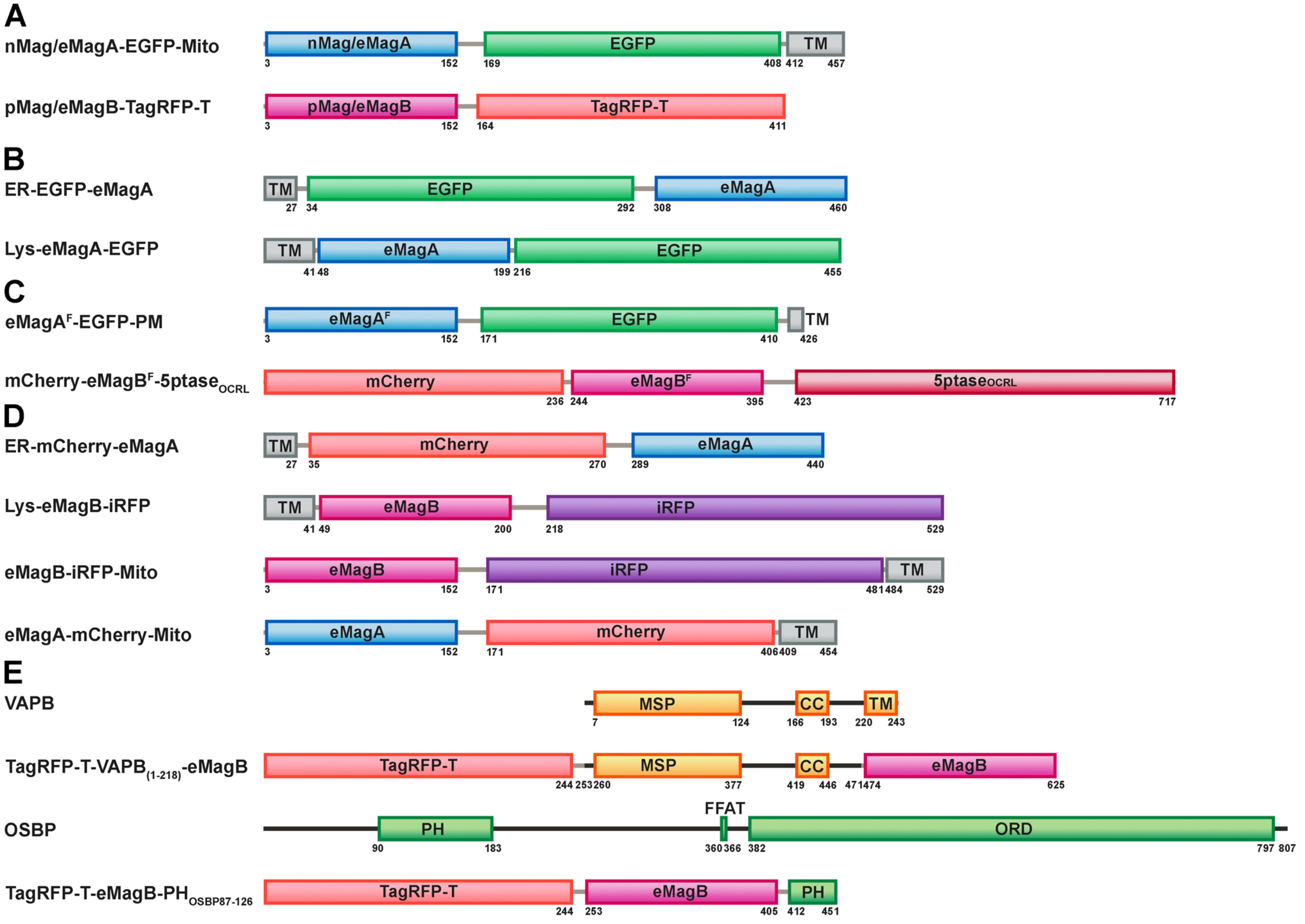

#### Supplementary Figure 2

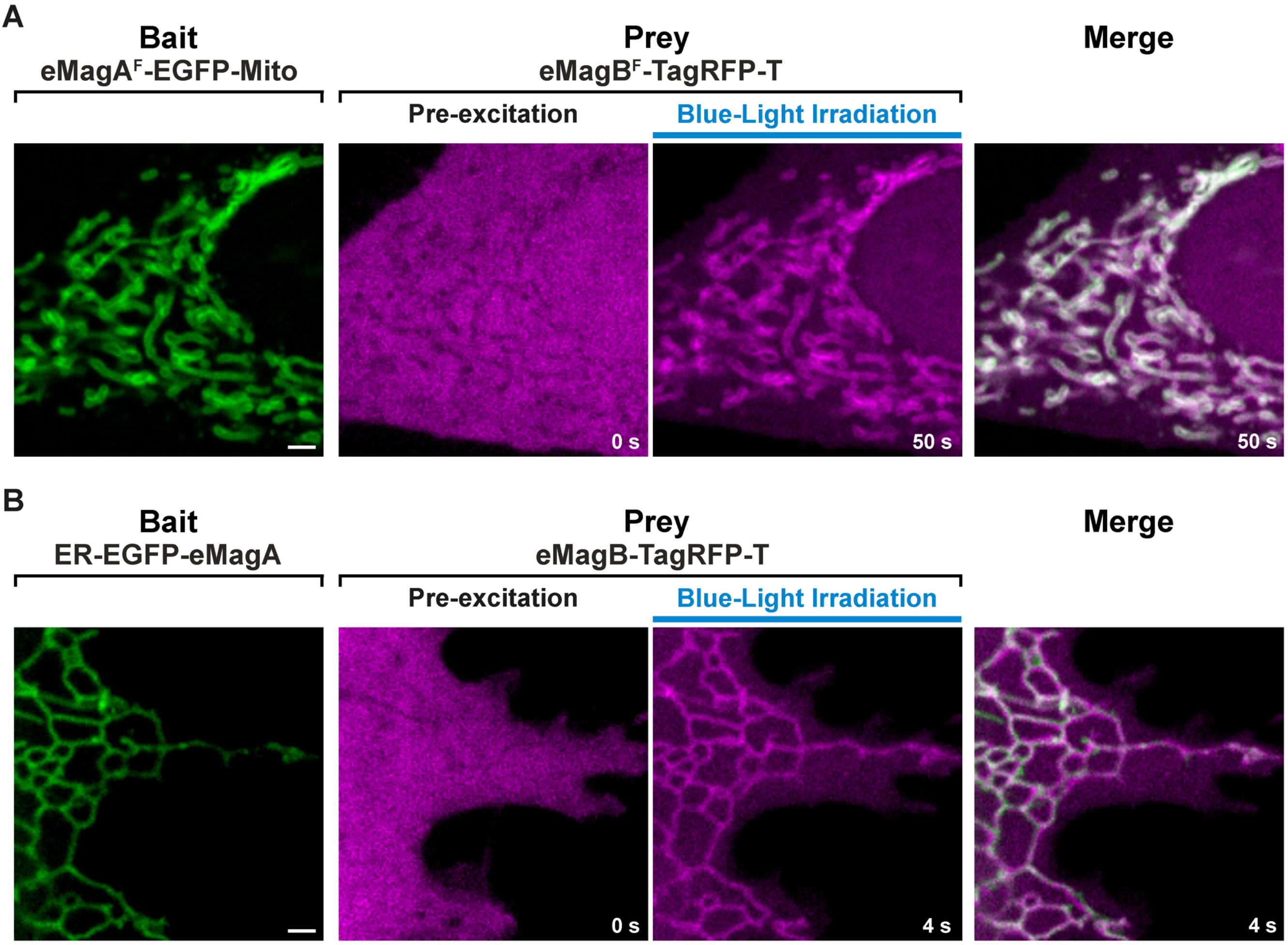

### Supplementary Figure 3

A

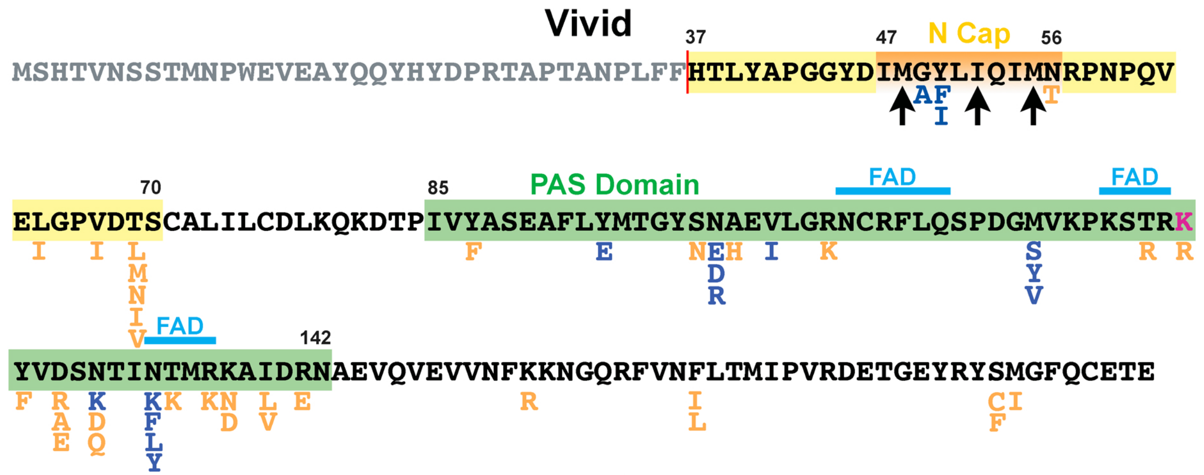


B

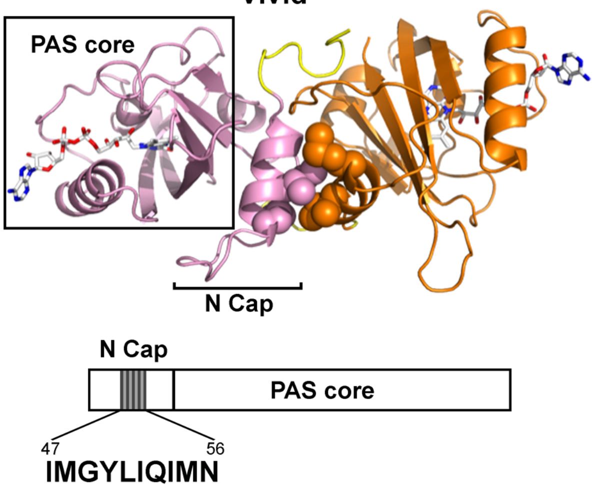


C

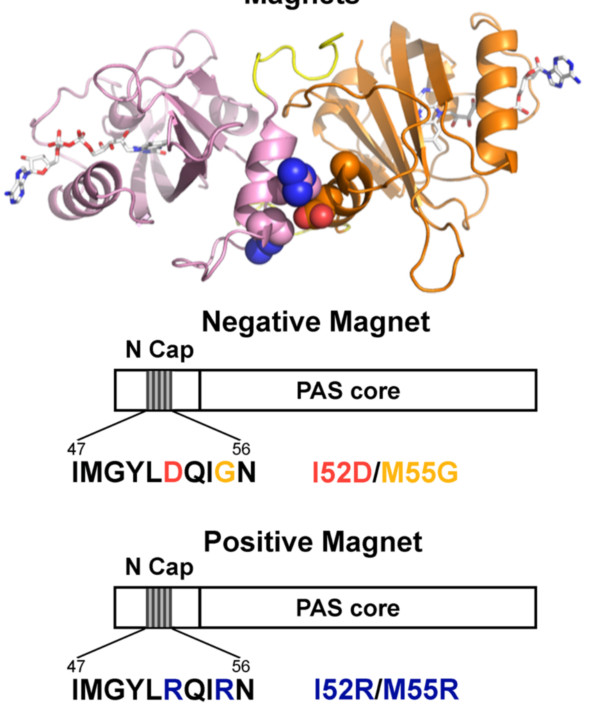


D

##### Alternative Complementary Charge Tested

| nMag | pMag | Result |
|---|---|---|
| I52E/M55G | I52R/M55R | Dimerization (-) |
| I52D/M55A | I52R/M55R | Dimerization (++) |
| I52D/M55D | I52R/M55R | Dimerization (+) |
| I52D/M55E | I52R/M55R | No dimerization |
| I52D/M55G | M48R/I52R/M55R | No dimerization |

**Supplementary Figure 4**

```
                                     55            69                       94
Wt                  LYAPGGYDIMGYLIQIMNRPNPQVELGPVDTSCALILCDLKQ-KDTPIVYASEAFLYMTGY
Thermomucor indicae LYTSTGLDVLSVLSRVVNRPNPEINVGPVDLSTAFLVVDAQAQFDFPIIYASPTFEQLTGY
Rhizomucor pusillus                ILLRPNPQINLGPIDMSCSFLVTDARQ-YDCPIVYCSPNFETLTGY
Rhizomucor miehei                       NPHINLGPVDFSCAFVVVDARQ-YDLPIAYVSPQFERLTGY
Thielavia           VYSKSGFDMLRALYYVATRKNPTVEIGAVDMSCSFIVCDLTL-NDCPIIYASDNFQNLTGY
Myceliophthora      VYSKSGFDMLRALFYVATRKNPTVEIGAVDLSCAFLVTDVTL-NDCPIIYVSDNFQNLTGY
Thermothelomyces    VYSKSGFDMLRALFYVATRKNPTVEIGAVDLSCAFLVTDVTL-NDCPIIYVSDNFQNLTGY
Chaetomium          IYSKSGFDMVRALAYVANRKNPTVEIGAVDFSCAFVVTDVTL-NDCPIIYVSDNFQNLTGY
other               IYSKSGFDMLRALWYVASRKDPKLKLGAVDMSCAFVVCDVTL-NDCPIIYVSDNFQNLTGY
                    :********:*** ***.**:*.:::****:**:*:* *:** *******.**********

                    99-101                   126    133  136
wt                  SNAEVLGRNCRFLQSPDGMVKPKSTRKYVDSNTINTMRKAIDRNAEVQVEVVNFKKNGQR
Thermomucor indicae PGREIVGRNCRFLQSPDGNVAQGSRRKFTDNNTVHVIRQDIIEGKETQSSLINYKRSGQP
Rhizomucor pusillus RNSEILGRNCRFLQAPDGQVTGGSRRQYTDNLAVYHLKSYLLQCKEHQASIINYRKGGQP
Rhizomucor miehei   SAREVIGRNCRFLQAPDGRVAIGSRRRYTDNSTAYHIKTHIIQGKESQCSIINYRKSGQ
Thielavia           NRHEIVGKNCRFLQSPDGVVEAGSRREFVANDAVFKLKNALAEGREIQQSLINYRKGGKP
Myceliophthora      NRHEIIGKNCRFLQSPDGEVEAGSRREFVANDAVLKLKNAVTEGKEIQQSLINYRKGGKP
Thermothelomyces    NRHEIIGKNCRFLQSPDGEVEAGSRREFVANDAVLKLKNAVTEGKEIQQSLINYRKGGKP
Chaetomium          NRHEIVGKNCRFLQSPSGVVEAGSRREFVANDAVFKLKNAVAEGKEIQQSLINYRKGGKP
other               SRHEIVGRNCRFLQAPDGNVEAGTKREFVENNAVYTLKKTIAEGQEIQQSLINYRKGGKP
                    .****:*:******:*.* ****::**** *:** .**::::**:***************

                                         179
wt                  FVNFLTMIPV-RDETGEYRYSMGFQCE
Thermomucor indicae FVNLLTVVPVLSKESNQIEYFVGFQID
Rhizomucor pusillus FVNLVTVIPI-QDDNEETAFFVGLQVD
Thielavia           FLNLLTLIPI-PWDNDKMKYCIGFQID
Myceliophthora      FLNLLTLIPI-PWDSDEIKYFIGFQID
Thermothelomyces    FLNLLTLIPI-PWDSDEIKYFIGFQID
Chaetomium          FLNLLTMIPI-PWETDEIKYCIGFQID
other               FLNLLTMIPI-PWDTEEIRYFIGFQID
                    ******:*** **:.::::* ******
```

**Supplementary Figure 5**

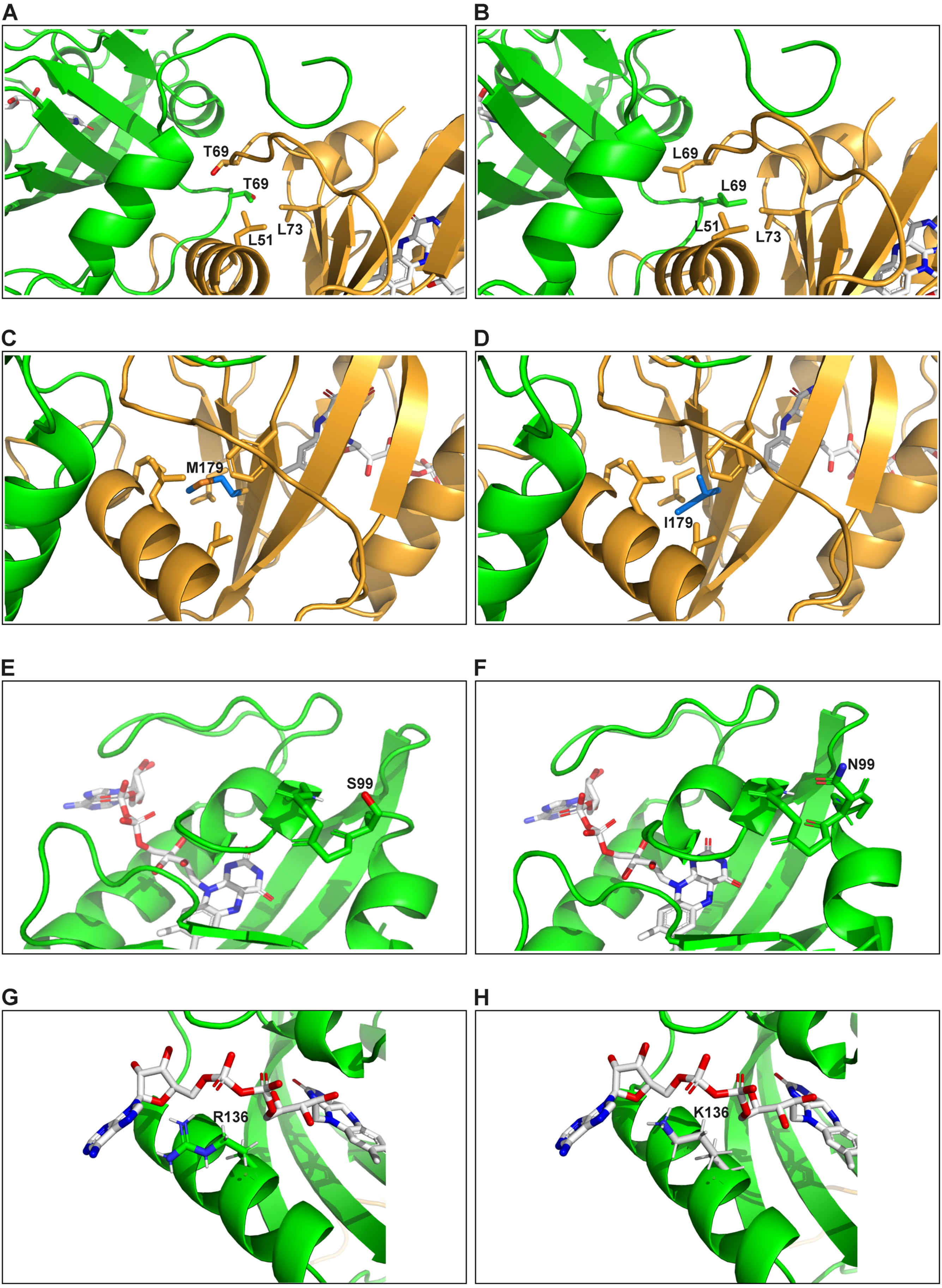

#### Supplementary Figure 6

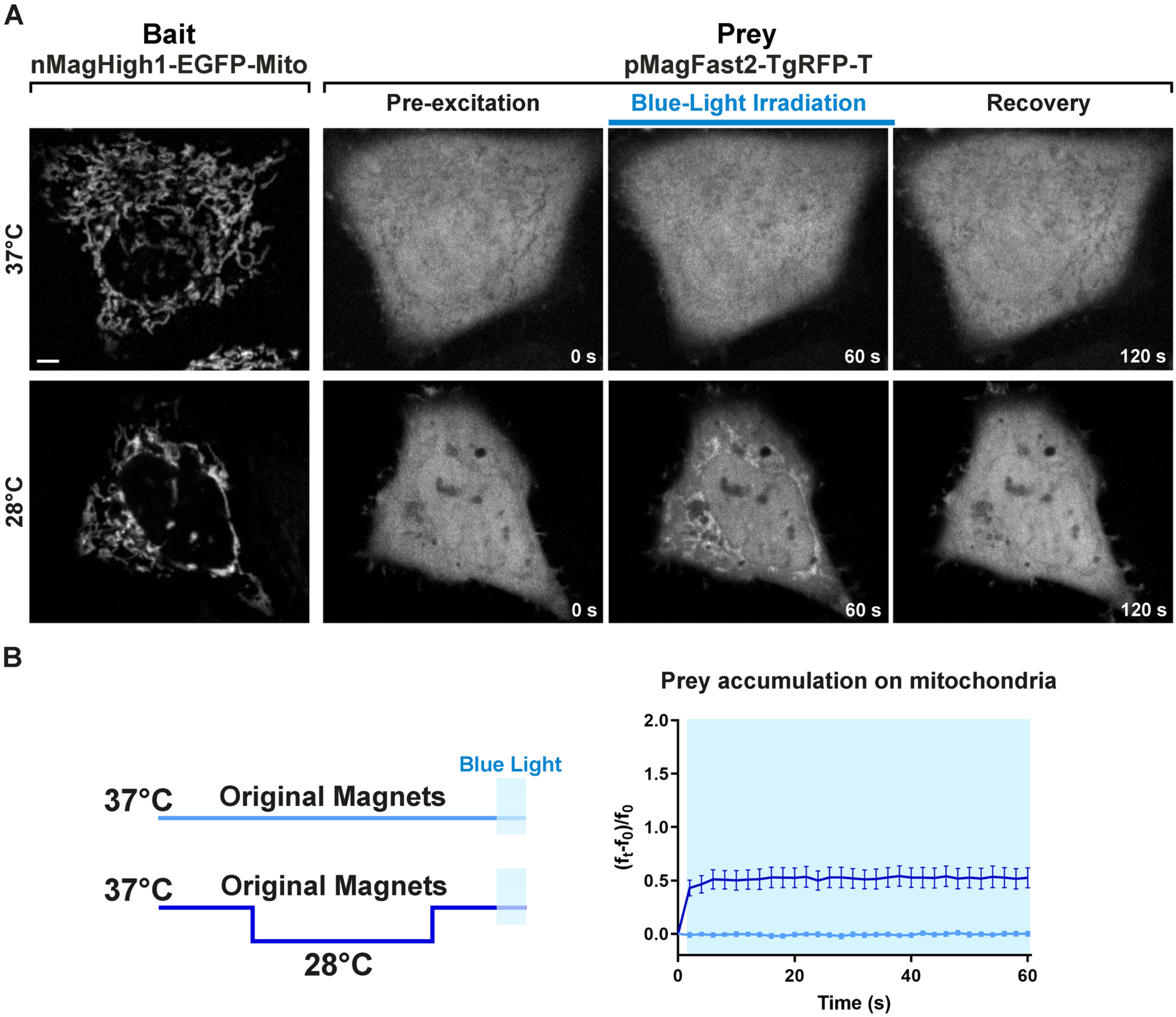

#### Supplementary Figure 7

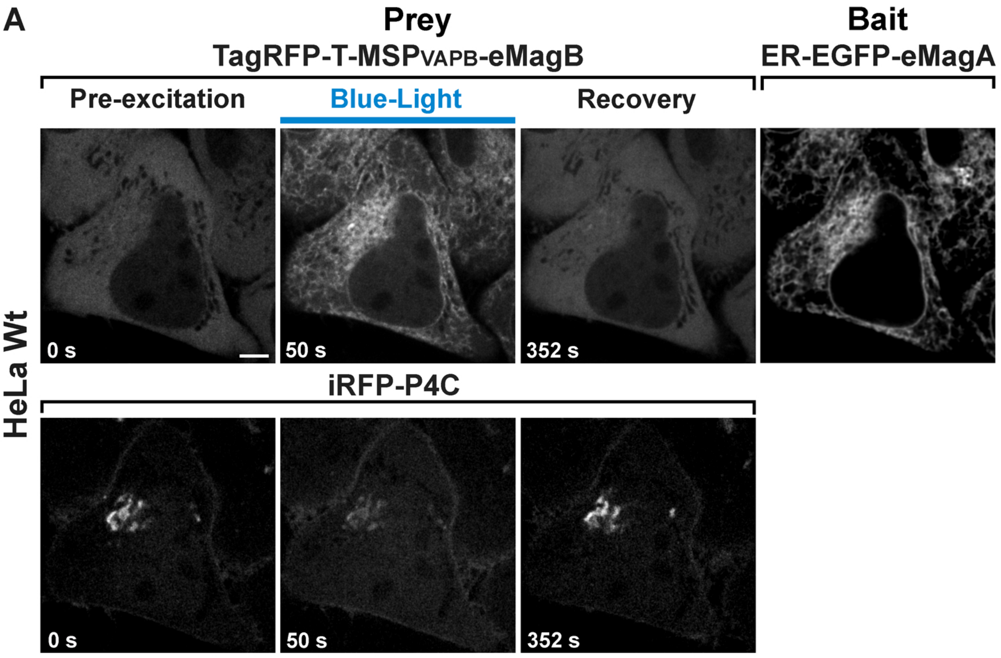


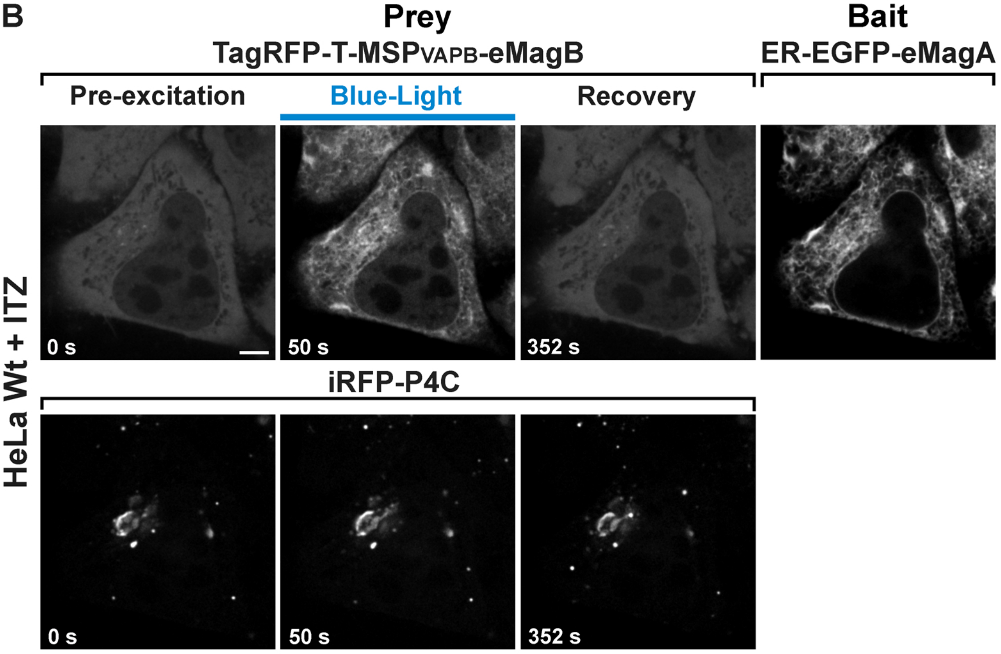

#### Supplementary Figure 8

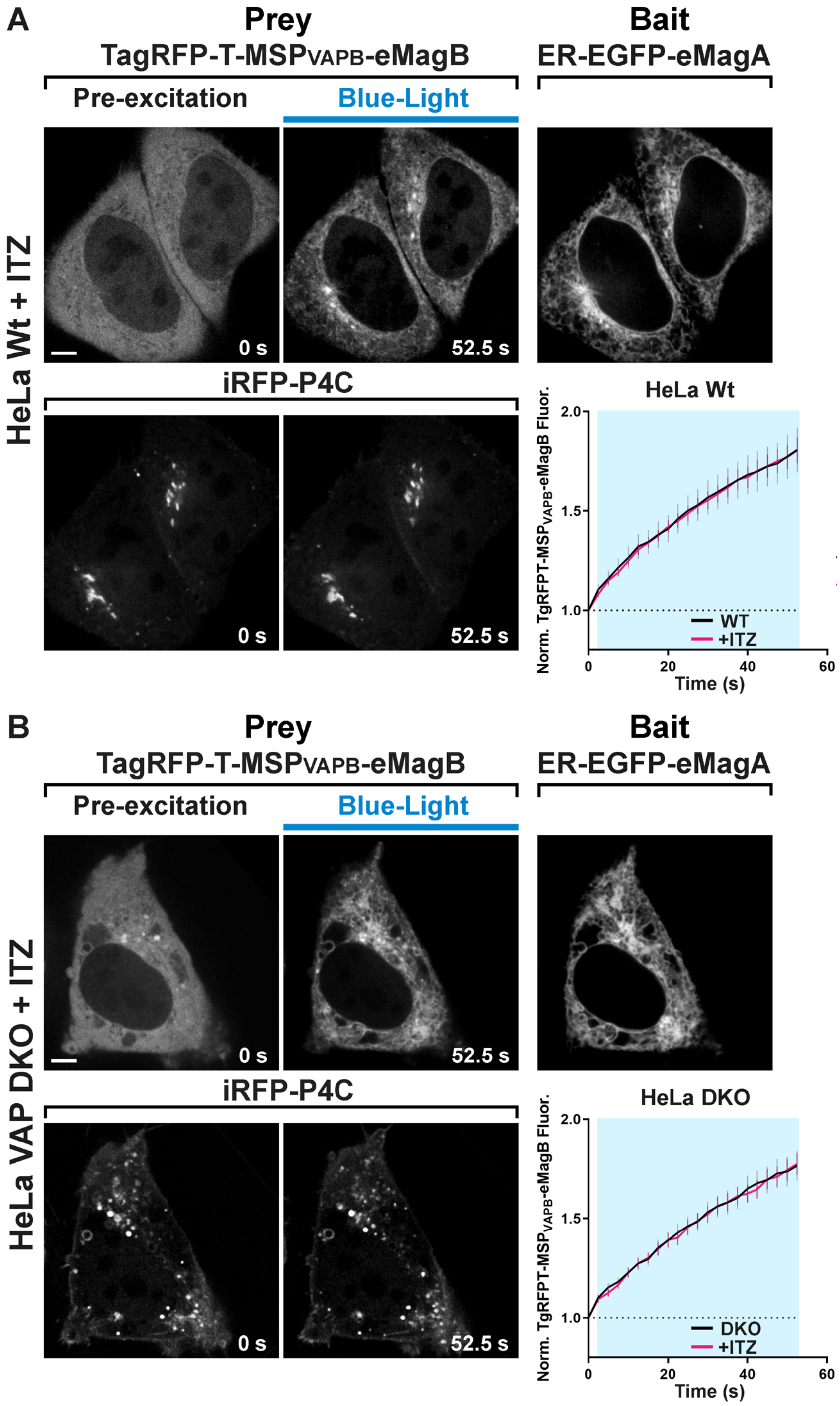

### Supplementary Figure 9

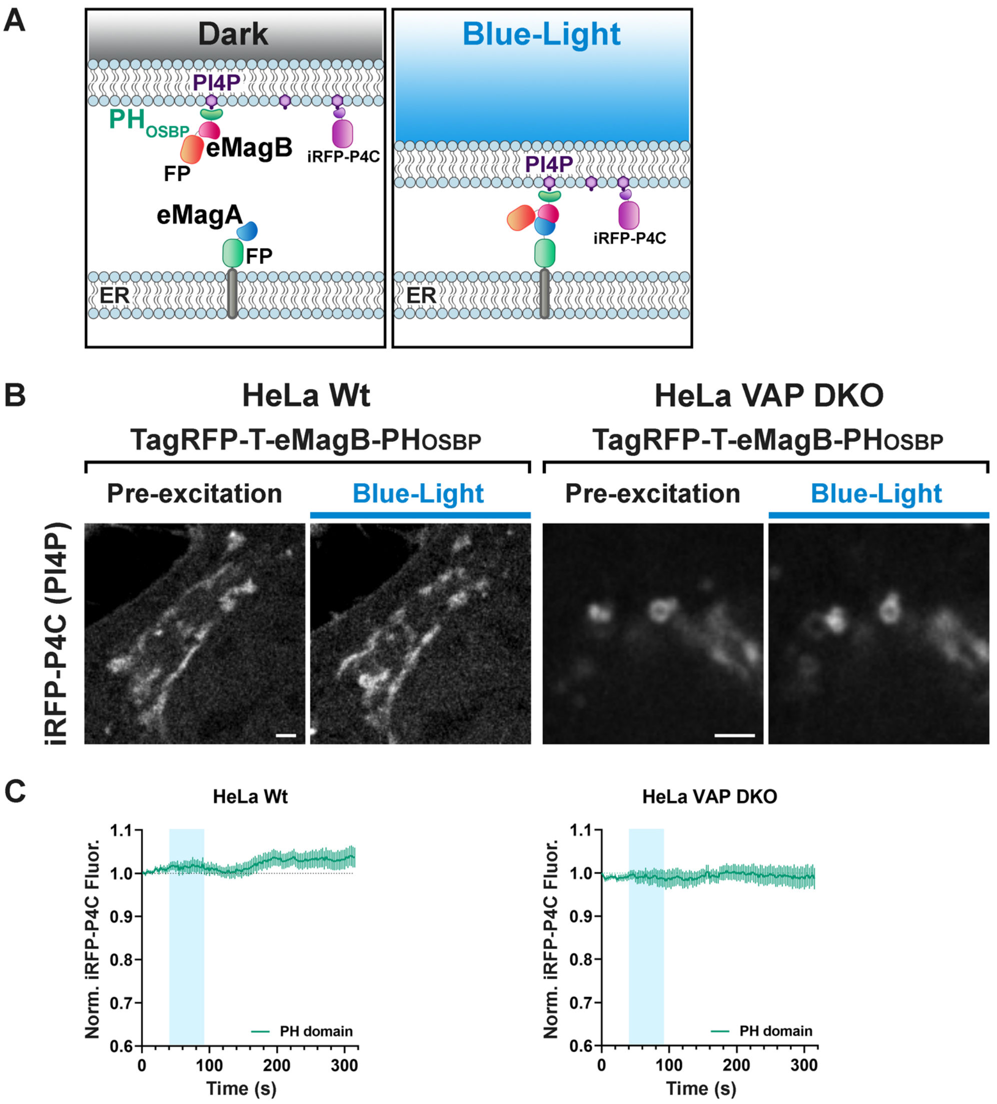
